# Prediction of Heart Disease and Survivability using Support Vector Machine and Naive Bayes Algorithm

**DOI:** 10.1101/2023.06.09.543776

**Authors:** Tanvi S. Patel, Daxesh P. Patel, Mallika Sanyal, Pranav S. Shrivastav

## Abstract

**Purpose:** In the present work, we examined the outcomes and accuracy of the Support vector machine (SVM) and the Naive Bayes algorithms on a dataset, to predict whether the patient has heart disease or not, and the patient’s survival prediction status.

**Method:** The machine learning procedures were developed using the clinically validated datasets with sixteen attributes from the University of California, Irvine’s Centre for Machine Learning, and Intelligent Systems. Confusion matrix was used to visualise the accuracy, recall, precision, and error of the models. Statistical analysis was done to prove the model accuracy using the receiver operating characteristic (ROC) curve and area under the curve (AUC).

**Results:** The proposed method of heart disease prediction using Naïve Bayes had 87 % accuracy. The accuracy for heart survivability models using SVM and Naïve Bayes were 88 % and 93 %. The model efficiency for heart survivability using ROC curve with AUC 0.93 for Naïve Bayes and AUC 0.91 for SVM.

**Conclusion:** Such prediction systems can help the medical sector to save energy, cost, and time by providing more efficient techniques to forecast decisions with high accuracy. This study will enable the statisticians and researchers to select more efficient and accurate machine learning algorithms to achieve better prediction of the “cardiovascular disease”.

## Introduction

Cardiovascular diseases are one of major causes of death worldwide. They are characterized as disorders of heart and blood vessels. This includes coronary heart disease, cerebrovascular disease, peripheral arterial disease, rheumatic heart disease, congenital heart disease and deep vein thrombosis and pulmonary embolism. According to the World Health Organization, approximately 18 million people die each year because of heart disease, accounting for nearly 31 % of all deaths [**1**]. Heart is a vital organ of the human body and with the aid of bioinformatics, hospitals, and non-governmental organizations (NGOs) are now using data to provide meaningful information. Applying data analytics to the health care domain will result in improved healthcare services. Data analytics is an emerging tool that is used in several sectors, such as healthcare, agriculture, energy, manufacturing, and many others [**2, 3**]. Data analytics helps to process the raw data, provide logical solutions and aid in decision making.

In the current world of technology, there are various types of smart phones and smart watches that can measure heart rate and rhythm and provide appropriate advice to keep the heart healthy. Machine learning (ML) is a branch of computer science that employs machine for solving patterns with large amounts of data and to assess possible results based on statistical analysis [**4**]. Various ML tools are being used to analyse several complex problems and offer possible solutions by constructing a model and identifying patterns in the data. ML focusses on development of intelligent algorithms that can help to analyse the fed data and provide a useful output. Mainly there are three type of ML algorithms namely, supervised learning, unsupervised learning, and reinforcement learning [**5**]. In the present work, we utilized the Support vector machine algorithm and the Naive Bayes algorithm, both of which are supervised algorithms. The work is basically on development of prediction model for heart disease and heart survivability. Multiple variables were used to predict the two outcomes. In the first estimation, there were 13 independent characteristics in the dataset as ‘heart-disease’ and 1 dependent result as ‘heart-disease-result*’*, and the prediction outcome was obtained utilizing these variables. In the second estimation, there were 12 independent characteristics as ‘heart-survivability’ and 1 dependent outcome as ‘heart-survivability-result’, and the prediction for survivability was generated utilizing these variables.

In the first experiment, Support vector machine (SVM) model and Naïve Bayes model were built on the training set ‘heart-disease’, ‘heart-disease-result’ and ‘heart-survivability’, ‘heart-survivability-result’ using the linear kernel. The entire outcome of the heart disease was obtained from 270 samples, 13 features variables and 1 target variable. In the second experiment, the same models were used for heart survivability data and to get an accurate prediction with the help of 299 samples, 12 features variables and 1 target variable. Further, a comparative assessment of both the model and algorithms is also provided with the existing work in the literature. The most prominent algorithm for classification issues is the Naive Bayes algorithm, which is also used to handle the multiclass problem. With independent variables, Naive Bayes performs better than logistic regression or other classification techniques. Both the algorithms were used in this statistical experiment to show which method performs better with the categorized data and that all variables in the dataset are assumed to be independent.

## Data sources, methodologies, and machine learning algorithms

### Data Source

In this prediction we have used the dataset from School of Information and Computer Science, University of California’s dataset called predict the heart disease and heart patient’s survivability [**6, 7**]. In the heart disease prediction, there are thirteen independent data attributes and one goal dependent attribute. The thirteen independent data attributes include age, sex, kind of chest pain, blood pressure, cholesterol, fasting blood sugar (FBS) over 120, electrocardiography (EKG) results, Max heart rate (HR), exercise angina, ST depression, ST slope, number of vessels Fluro (number of major vessels (0-3) coloured by fluoroscopy), and thallium (**Table 1)**. There are twelve independent data attributes and one target dependent attribute in the heart failure clinical records dataset. Age, anaemia, creatinine phosphokinase, diabetes, ejection fraction, high blood pressure, platelets, serum creatinine, serum sodium, sex, smoking, and time are the twelve independent data attributes and one dependent target attribute death_event as presented in **Table 2**. The analysis of data features and prediction results code is available on the GitHub account [**8**].

**Table 1.**
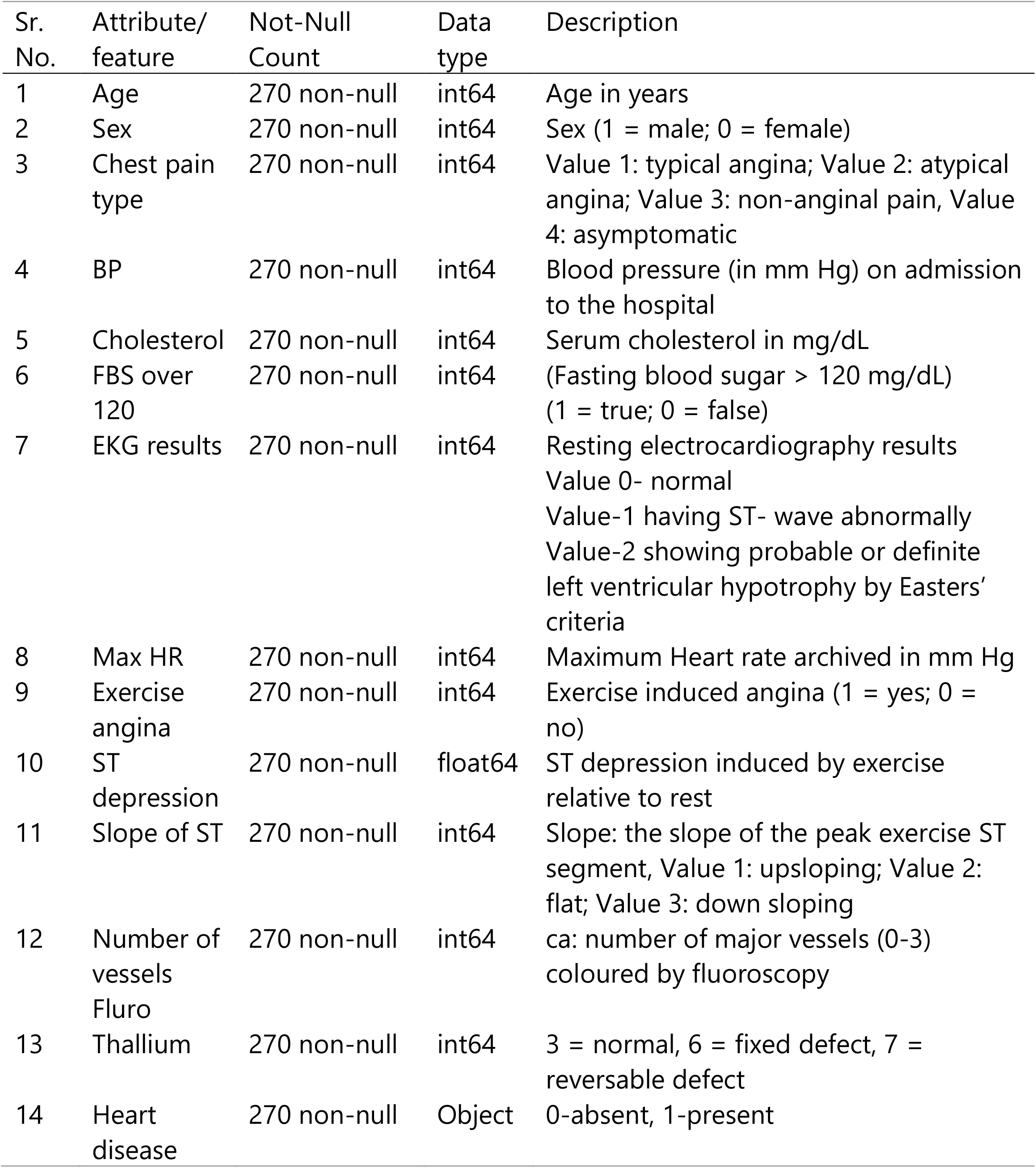
Heart disease prediction dataset

**Table 2.** Heart survivability dataset

| Sr. No. | Attribute/feature | Not-Null Count | Data type | Description |
| --- | --- | --- | --- | --- |
| 1 | Age | 299 non-null | float64 | Age in years |
| 2 | Anaemia | 299 non-null | int64 | Decrease of red blood cells or haemoglobin (1=yes, 0=no) |
| 3 | Creatinine_phosphokinase | 299 non-null | int64 | Level of the CPK enzyme in the blood (mcg/L) |
| 4 | Diabetes | 299 non-null | int64 | If the patient has diabetes (1=yes, 0=no) |
| 5 | Ejection_fraction | 299 non-null | int64 | Percentage of blood leaving the heart at each contraction (percentage) |
| 6 | High_blood_pressure | 299 non-null | int64 | If the patient has hypertension (1=yes, 0=no) |
| 7 | Platelets | 299 non-null | float64 | Platelets in the blood (Kilo platelets/mL) |
| 8 | Serum_creatinine | 299 non-null | float64 | Level of serum creatinine in the blood (mg/dL) |
| 9 | Serum_sodium | 299 non-null | int64 | Level of serum sodium in the blood (meq/L) |
| 10 | Sex | 299 non-null | int64 | Woman or man |
| 11 | Smoking | 299 non-null | int64 | If the patient smokes or not (1=yes-, 0=no) |
| 12 | Time | 299 non-null | int64 | Follow-up period (days) |
| 13 | Death_event | 299 non-null | int64 | If the patient is deceased during the follow-up period (1=yes, 0=no) |

### Flow chart for constructing prediction model

In this project, we used two categorized clinical datasets to predict two outcomes from the UCI machine learning repository, employing two machine learning methods (SVM and Naive Bayes). **Fig. 1** illustrates the process utilized to obtain correct results. First, the dataset was imported into the text editor to process it further and then separated the dataset into independent characteristic and a target binary value as a dependent variable. To facilitate the process, the binary variables were encoded. The first dataset shape had 14 attributes and 270 samples of the normal individuals, while in the second dataset there were 13 attributes and 299 samples of the heart patients. There were no missing values in both the datasets. The independent attributes were pre-processed for better outcome. After successful data pre-processing, it was split into training set and test set. Standard scaling feature was applied on the train and test set.

**Figure 1.**
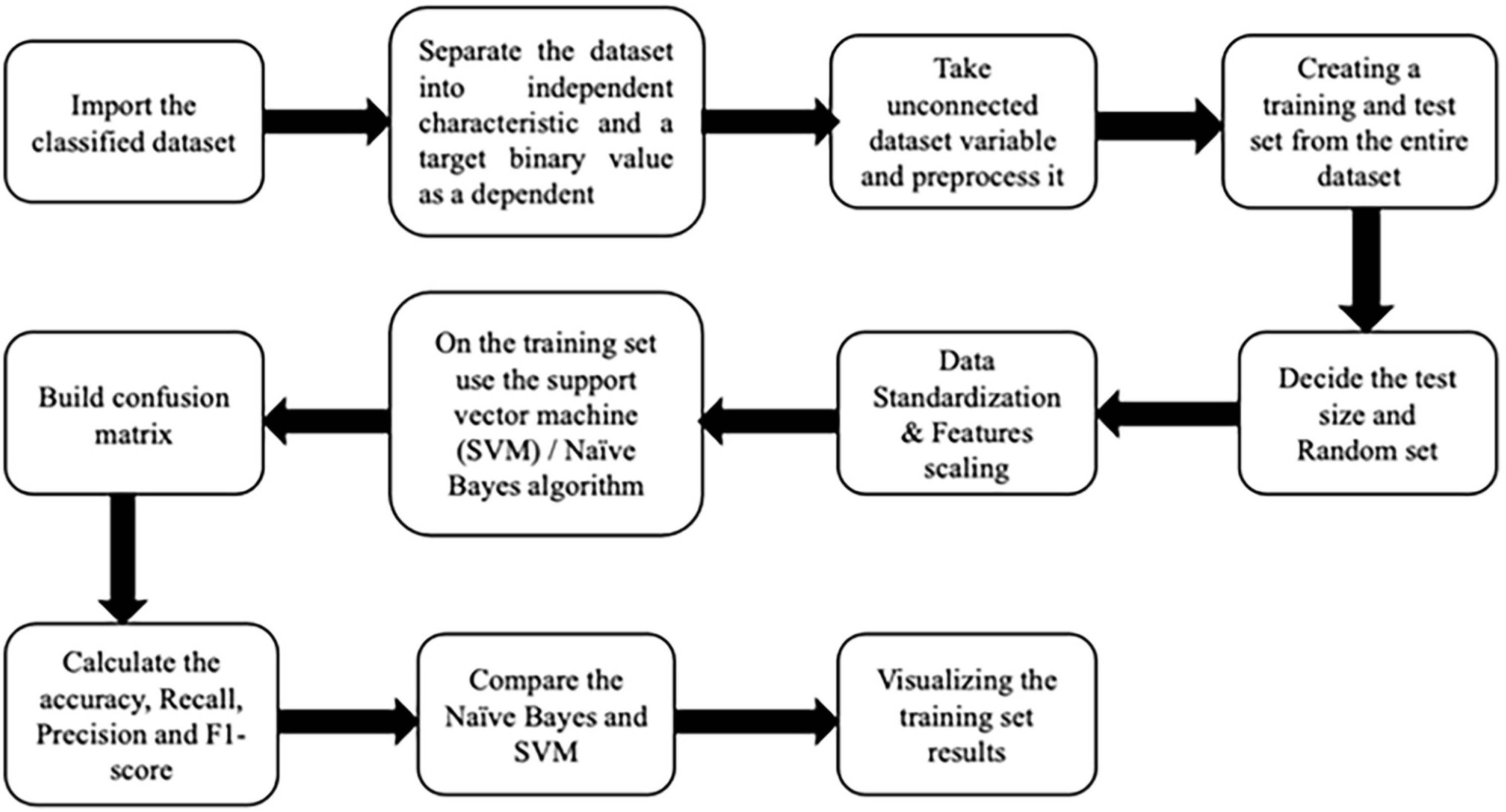
Flowchart for constructing the prediction model.

Equation for standard feature scaling: 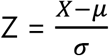

Here, x = value of X, *μ = mean value of X, σ = standard deviation of X*

Subsequently, SVM and Naïve Bayes algorithms were applied on the training set and predicted the test set results. Further, confusion matrix was applied to calculate the precision, recall, F1-score, and accuracy of the model. After completing the process on both datasets, the algorithm’s outcome and prediction were compared and the results were visualized using various graphs.

### Support vector machine

SVM methodology was utilized to determine the pattern of the dataset to predict heart disease and cardiac survival. SVM was created by AT&T Bell Labs’ Vladimir Vapnik. It is based on the concept of decision planes, which are used to draw decision lines [**9, 10**]. SVM can be used to solve both regression and classification problems, albeit we only utilized it to address the classification problem in this case. In this experiment, we chose the quadratic SVM model, which is also operate well when data is misclassified and not around the hyperplane.

### Kernel SVM

The essential technology in SVM is the kernel function; different kernels lead to different SVM classifications [**11**]. The linear indivisible data of space map to high dimension nonlinear space via the kernel function, enables sample linear classification in high dimension space, namely, the inner product operation of all vectors in high dimension space via the kernel function of original space. There are many types of kernels used in SVM algorithm like polynomial kernel, linear kernel, RBF kernel function, sigmoid kernel function and others. In this experiment we have used linear kernel function for the proper dimensions of the model. The mathematical equation used for the linear kernel was

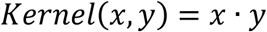

### SVM equation to solve the classification problem

The problem of supervised learning is formulated as follows in statistical learning theory. We are given a set of training data (*x*1, *y*1)… (*x*l, *y*l) in Rn R sampled according to an unknown probability distribution P(*x, y*), as well as a loss function V(*y*, f(*x*)) that measures the variance, where f(*x*) is “predicted” instead of the actual value ‘*y*’ for a given ‘*x*’ [**12**]. In this classification, the best hyperplane is where the margin is maximum.

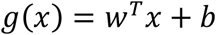

In the linear graph, g(x) is *w*^*T*^*x + b* where b is width, ‘*x*’ is input and w is the separator.

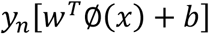

Result is negative if the given equation is less than zero and result is positive if the given equation is greater or equal to zero.

### Naïve Bayes algorithm

Naïve Bayes is the algorithm used for different attributes of the heart disease dataset to predict the probability of the disease. The Bayes theorem allows to calculate the posterior probability, P(c|*x*), from P(c), P(*x*), and P(x|c) using P(c), P(x), and P(x|c). The effect of the value of a predictor (*x*) on a given class (c) is unrelated to the presence of other predictors, according to the Naive Bayes classifier. This is known as class conditional independence [**13**]. There were 13 features and 1 class in the first experiment, each of which is dependent on 13 independent attributes. In the second experiment, there were 12 independent traits and 1 class that relies on the 12 features for accurate results. In Naive Bayes, all independent attributes are entered as ‘*x*’ and output as class ‘*c*’ or ‘*y*’, and the algorithms provide accurate results using this pattern.

### Naïve Bayes to solve the classification problem

Bayes theorem shows the relationship between the independent features and dependent outcomes and is expressed as,

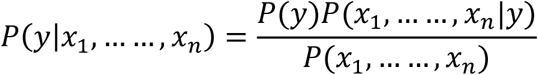

Consider

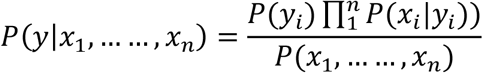

where 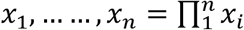 is the probability of ‘*x*’ occurring when ‘*y’* has already occurred, *P*(*x*_1,….,_*x*_*n*_) is the probability of ‘*x*’ to occur; *P*(*y*_*i*_) is the probability of ‘*y’* to occur, and *P*(*y*|*x*_1_, … …, *x*_*n*_) is the probability of ‘*y*’ occurring when ‘*x*’ is already occurred **[14]**.

## Statistical analysis

### Various attributes of the dataset with dependent values

As there are many changes in the individual features to dependent values, median and mean are not adequate to estimate the dataset’s probability. Thus, we utilized violin plots in this experiment to demonstrate the range of the different independent attributes to dependent attributes. Different characteristics of violin plots of the heart disease datasets are displayed in **Fig. 2a**. The ST-depression violin plot shows that the biggest probability of heart disease is between values 1 and 2. In patients with heart disease, the median value of ST depression is greater than in healthy people. Among the dataset, the maximum heart rate is lower in heart patients, and most patients near the age of 60 are heart patients. As evident from the figure, the individual with chest pain has the highest likelihood of having heart disease. Besides, it is difficult to predict the heart disease just with blood pressure and cholesterol levels. **Fig. 2b** depicts the likelihood of death as a function of serum sodium, serum creatinine level, time (days) of heart disease, and ejection fraction. The patients’ chances of surviving their heart attacks are increasing with time. In the time violin plot, the chance of a death event is highest when the number is less than 50. The amount of serum creatinine is also critical in assessing the likelihood of mortality. In terms of serum creatinine, the median is greater than the average individual.

**Figure 2.**
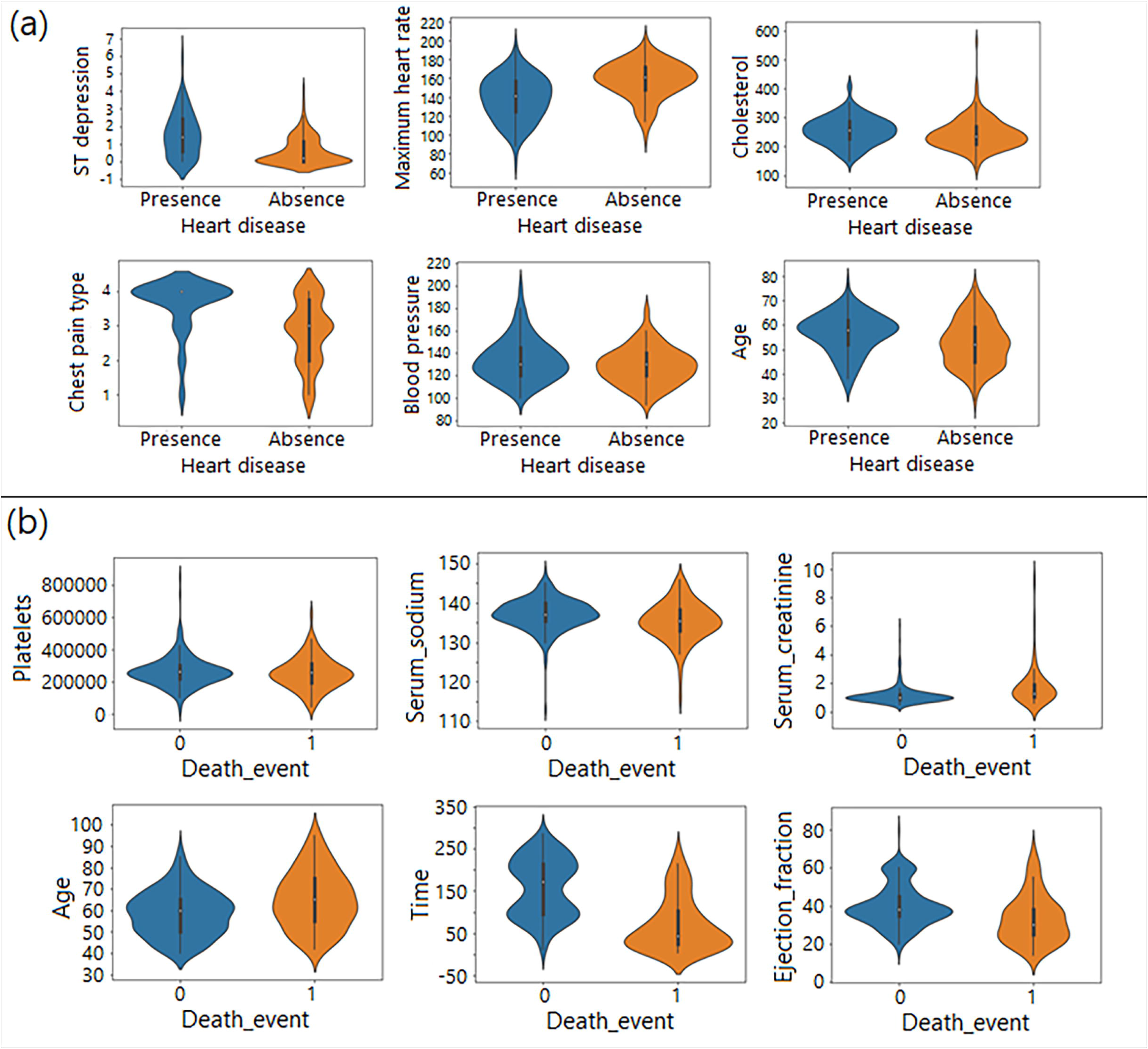
Violin plots for (a) heart disease data and (b) heart survivability data

### Importance of attributes and feature selection

The dataset contains many features and attributes, some of which are significant while others are insignificant. If there are some non-important attributes, then to improve model accuracy and prediction, feature selection approach can be used [**15**]. In some cases, unused attributes take longer to forecast in the model. Another application of feature selection is to determine the significance of characteristics in the target values. In this experiment, the model was trained with all the dataset properties, as they are all important factors in predicting heart disease and survival. Besides, hospitals typically assess all the primary needs of patients and therefore the experiment considered all the attributes of the dataset. This experiment used the information gain technique to measure the importance of the variables. The decrease in entropy caused by a dataset transformation is measured as information gain. It can be used to choose features by evaluating each variable’s information gain in relation to the target variable. **Fig. 3** defines the importance of independent dataset features in archiving goal values and in model construction, as well as the attributes utilized in which amounts. In **Fig. 3a**, the chest pain type, ST depression, thallium, and number of vessels Fluro are the important attributes of heart disease, while serum sodium, serum creatinine and time are important attributes of the heart survivability (**Fig. 3b**). **Confusion matrix**

**Figure 3.**
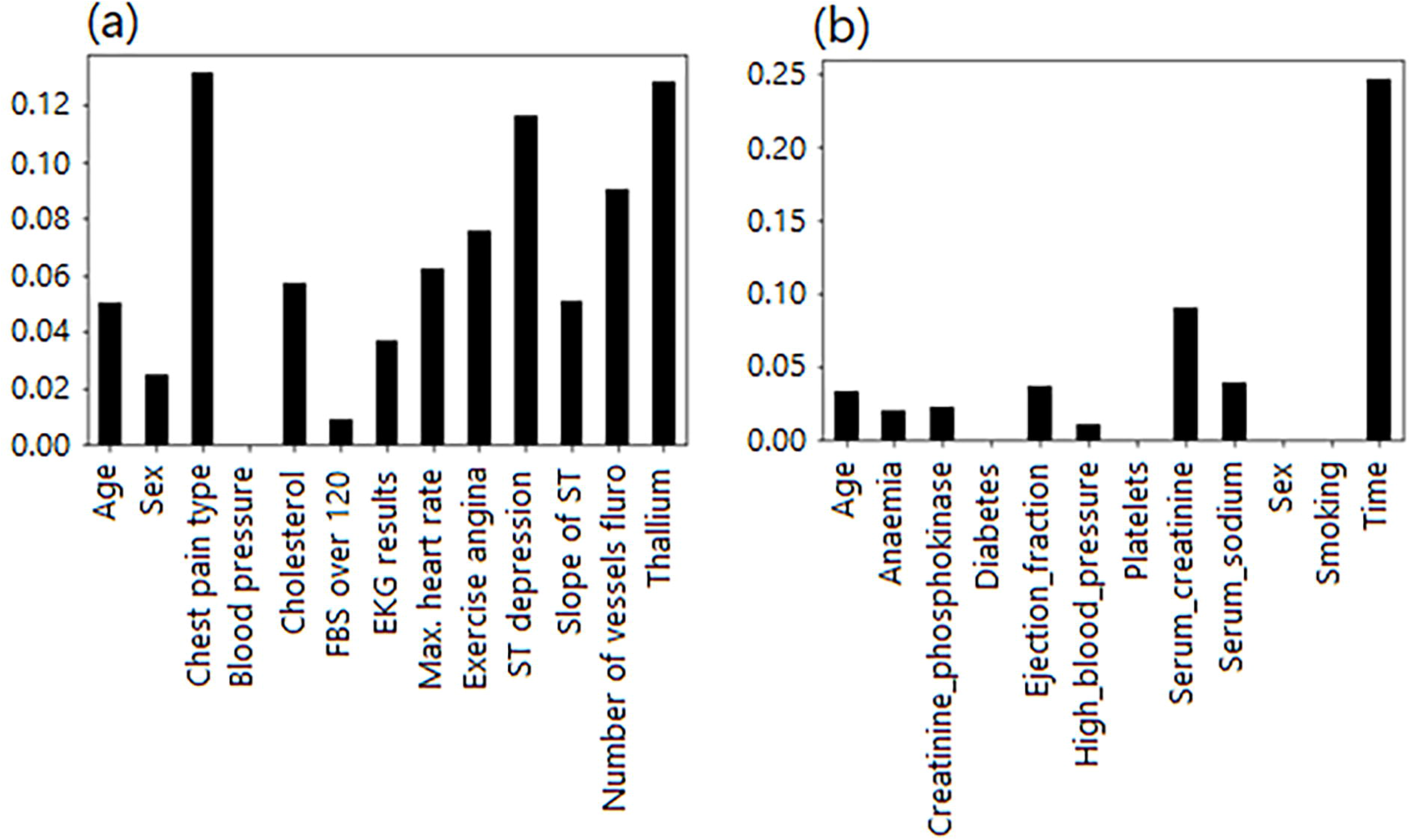
Measured features statistics obtained from the prediction model for (a) heart disease and (b) heart survivability.

In this experiment we have used the confusion matrix to visualise the accuracy, recall, precision, and error of the models. The anticipated and actual classification are shown in a confusion matrix of dimension *n × n* connected with a classifier, where *n* is the number of distinct classes [**16, 17**]. In this experiment the value of *n* is 2 because we have used 2 classes in four of the machine learning models.

Accuracy is defined as the ratio of properly predicted occurrences to all instances in the dataset [**18**]. Precision is also known as predicted positive occurrences so basically precision is the ratio of true positive predicted occurrences to true predicted occurrences. Whereas recall is the ratio of true positive predicted occurrences to true positive predicted occurrences plus false negative predicted occurrences. The F1 score represents the balance between precision and recall [**19**]. **Fig. 4** shows the confusion matrixes for all the models with true positive, false positive, true negative and false negative numbers. In this figure, the TPN, FPN, TNN, FNN represent true positive predicted numbers, false positive predicted numbers, true negative predicted numbers, and false negative predicted numbers, respectively.

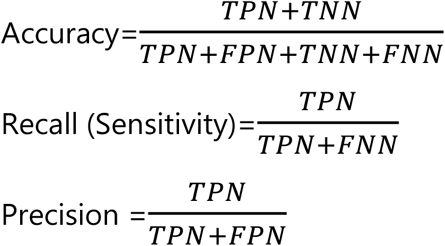

To find the accuracy, precision, recall in the models, the following equations and confusion matrix numbers were used.

**Figure 4.**
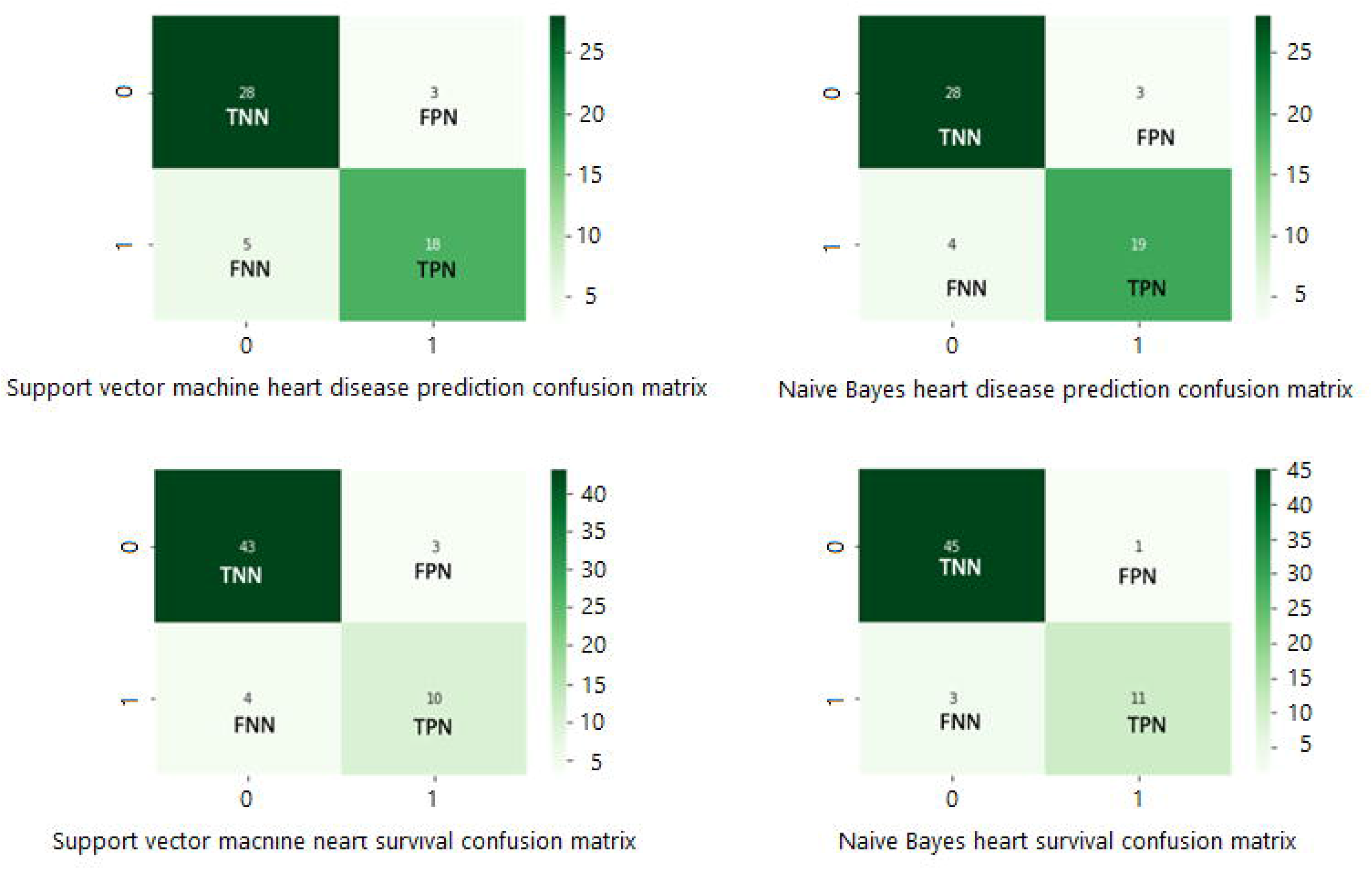
Experiment of confusion matrices to calculate the results.

## Results and discussion

The purpose of this study was to achieve higher accuracy in the heart prediction as well as heart survivability as compared to previous studies using the Naïve Bayes and SVM algorithm. As several patients who have heart ailments are susceptible to heart stroke it is useful to have a model which could predict any such event early and foresee patient’s survivability using the same datasets. The data in **Table 3** shows the method accuracy, error, precision, and recall values using SVM and Naïve Bayes algorithms. The proposed method of heart disease prediction using Naïve Bayes had 87 % accuracy, which is higher as compared to previous reports [**4, 7, 9, 16, 20**]. **Table 4** shows a comparison of method accuracy using Naïve Bayes and SVM algorithms with reported work. The proposed method of heart disease prediction using SVM also had an accuracy of 85%, which is better as compared existing reports. Heart survivability methods are unique and relatively new and as such there are few reports on methods developed using these tools and dataset. The accuracy for heart survivability models using SVM and Naïve Bayes were 88 % and 93 %, which is higher than one report [**4**], that used the same dataset to make the prediction model.

**Table 3.** Model performance with 80 and 20 percent of test size

| Model | Accuracy | Error | Precision | Recall | AUC |
| --- | --- | --- | --- | --- | --- |
| HDP using SVM | 0.85 | 0.15 | 0.86 | 0.78 | 0.87 |
| HDP using Naïve Bayes | 0.87 | 0.13 | 0.86 | 0.83 | 0.89 |
| HSP using SVM | 0.88 | 0.12 | 0.77 | 0.71 | 0.91 |
| HSP using Naïve Bayes | 0.93 | 0.7 | 0.92 | 0.79 | 0.93 |

HDP: Heart disease prediction; HSP: Heart survivability prediction; SVM: Support vector machine

**Table 4.** Comparison of model accuracy with reported methods

| Model | Accuracy |  |  |  |  |  |  |
| --- | --- | --- | --- | --- | --- | --- | --- |
|  | Present method | Ref 7 | Ref 9 | Ref 16 | Ref 20 | Ref 4 | Ref 22 |
| HDP using SVM | 0.85 | 0.82 | 0.85 | 0.86 | <0.85 | -- | 0.82 |
| HDP using Naïve Bayes | 0.87 | 0.84 | 0.86 | 0.76 | <0.85 | -- | -- |
| HSP using SVM | 0.88 | -- | -- | -- | -- | 0.68 | -- |
| HSP using Naïve Bayes | 0.93 | -- | -- | -- | -- | 0.69 | -- |

HDP: Heart disease prediction; HSP: Heart survivability prediction; SVM: Support vector machine

In the present model, statistical analysis was done to prove the model accuracy using the receiver operating characteristic (ROC) curve and area under the curve (AUC) [**21**]. Graphically it can be seen that for heart prediction model, AUC is 0.89 for the Naïve Bayes and 0.87 for SVM (**Fig. 5a). Fig. 5b** also shows the model efficiency for heart survivability using ROC curve with AUC 0.93 for Naïve Bayes model and AUC 0.91 for SVM. One reported method [**7**] evaluated the performance of six different machine learning techniques for prediction of heart disease for all attributes on datasets, however, their accuracy was slightly lower than our work. Mohan et al. [**16**], compared their proposed Hybrid Random Forest Linear Model (HRFLM) with several other machine learning algorithms. The obtained accuracy with Naïve Bayes was 0.76 and with SVM is 0.86. In another work [**9**], the accuracy using Naïve Bayes and SVM were 0.86 and 0.85. respectively, which are close to our work. A survey paper [**20**] suggests that when applying algorithms to all the attributes, the accuracy of Naïve Bayes is the highest at 84 %, and the accuracy of SVM at 85 %, indicating that the accuracy of this proposed method is slightly better. A prediction using supervised learning classifier **[22]** applied Cascaded Neural Network (CNN) and SVM to predict heart disease. They used a non-linear kernel function and the accuracy of the SVM model was 82 %, signifying that the present SVM approach is more accurate. Likewise, SVM with different datasets and characteristics was used to predict heart failure with an accuracy of 77 % **[23]**, further demonstrating the better accuracy of the proposed SVM method for heart survivability. Naive Bayes is used specifically for classification problems and considers all features to be independent, whereas SVM is used for both regression and classification problems. Nevertheless, Naive Bayes works better for categorised datasets than SVM.

**Figure 5.**
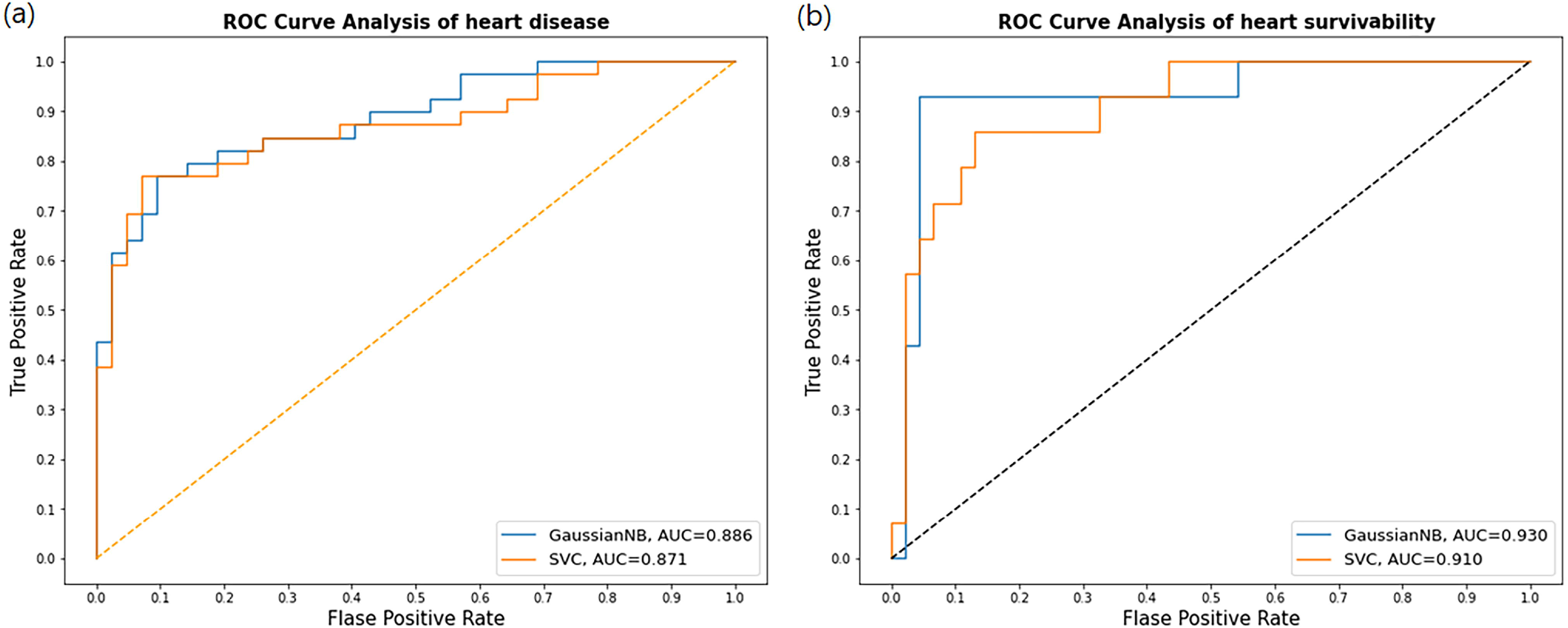
Receiver operating characteristics (ROC) curve for comparison of support vector machine and Naïve Bayes algorithms for (a) heart disease and (b) heart survivability.

## Conclusion

As heart disease and stroke are the major causes of death it has become essential to use technology to detect these outbreaks early and save human lives. The heart survivability model was shown to be 93 % accurate using Naïve Bayes in this study. In comparison to the SVM method, it has become evident that the Naïve Bayes algorithm performs best in terms of categorized prediction models. In comparison to previous research on same datasets that used split data in training and testing sets as well as run accurately, this research contributes higher accuracy.

## Declarations and Statements

## Acknowledgments

We would like to thank Department of Computer Science, Bowie State University and Department of Chemistry, Gujarat University for supporting this work.

## Author’s contributions

TSP devised and designed the study, wrote the study protocol, contributed to data collection and analyses.

DPP contributed to the study idea and protocol and did analyses.

MS contributed substantially towards writing the draft manuscript.

PSS contributed towards critically revising, editing the manuscript and supervision.

Further, all authors read and approved the final manuscript for submission

## Availability of data and materials

The data features and prediction results code are available on the GitHub account [8].

